# Refining epigenetic prediction of chronological and biological age

**DOI:** 10.1101/2022.09.08.507115

**Authors:** Elena Bernabeu, Daniel L McCartney, Danni A Gadd, Robert F Hillary, Ake T Lu, Lee Murphy, Nicola Wrobel, Archie Campbell, Sarah E Harris, David Liewald, Caroline Hayward, Cathie Sudlow, Simon R Cox, Kathryn L Evans, Steve Horvath, Andrew M McIntosh, Matthew R Robinson, Catalina A Vallejos, Riccardo E Marioni

## Abstract

Epigenetic clocks can track both chronological age (cAge) and biological age (bAge). The latter is typically defined by physiological biomarkers and risk of adverse health outcomes, including all-cause mortality. As cohort sample sizes increase, estimates of cAge and bAge become more precise. Here, we aim to refine predictors and improve understanding of the epigenomic architecture of cAge and bAge. First, we perform large-scale (N = 18,413) epigenome-wide association studies (EWAS) of chronological age and all-cause mortality. Next, to improve cAge prediction, we use methylation data from 24,673 participants from the Generation Scotland (GS) study, the Lothian Birth Cohorts (LBC) of 1921 and 1936 and 8 publicly available datasets. Through the inclusion of linear and non-linear age-CpG associations from the EWAS, feature pre-selection/dimensionality reduction in advance of elastic net regression, and a leave-one-cohort-out (LOCO) cross validation framework, we arrive at an improved cAge predictor (median absolute error = 2.3 years across 10 cohorts). In addition, we train a predictor of bAge on 1,214 all-cause mortality events in GS, based on epigenetic surrogates for 109 plasma proteins and the 8 component parts of GrimAge, the current best epigenetic predictor of all-cause mortality. We test this predictor in four external cohorts (LBC1921, LBC1936, the Framingham Heart Study and the Women’s Health Initiative study) where it outperforms GrimAge in its association to survival (HR_GrimAge_ = 1.47 [1.40, 1.54] with *p* = 1.08 × 10^−52^, and HR_bAge_ = 1.52 [1.44, 1.59] with *p* = 2.20 × 10^−60^). Finally, we introduce MethylBrowsR, an online tool to visualize epigenome-wide CpG-age associations.

## Introduction

The development and application of epigenetic predictors for healthcare research has grown dramatically over the last decade^1^. These predictors can aid disease risk stratification, and are based on associations between CpG DNA methylation (DNAm) and age, health, and lifestyle outcomes. DNAm is dynamic, tissue-specific and is influenced by both genetic and environmental factors. DNAm can precisely track ageing through predictors termed “epigenetic clocks”^2–8^. DNAm scores have also been found to capture other components of health, such as smoking status^9,10^, alcohol consumption^11,12^, obesity^11,13^, and protein levels^14^.

“First generation” epigenetic ageing clocks, including those by Horvath^3^ and Hannum et al^4^, were trained on chronological age^2–4^ (cAge), with near-perfect clocks expected to arise as sample sizes grow^5^. However, cAge clocks hold limited capability for tracking and quantifying age-related health status, also termed biological age (bAge)^5,8^. To address this, “second generation” clocks have been trained on other age-related measures, including a phenotypic biomarker of morbidity (PhenoAge^15^), rate of ageing (DunedinPoAm^16^), and time to all-cause mortality (GrimAge^17^). Regressing an epigenetic clock predictor (whether trained on cAge or bAge) on chronological age within a cohort gives rise to an “age acceleration” residual with positive values corresponding to faster biological ageing.

Penalised regression approaches such as elastic net^18^ are used to derive epigenetic predictors. Such epigenetic clocks typically capture a weighted linear combination of CpGs that optimally predict an outcome from a statistical perspective i.e. no preference is given to the location or possible biological role of the input features. The majority consider genome-wide CpG sites as potential predictive features. However, others have used a two-stage approach that first creates DNAm surrogates (or epigenetic scores - EpiScores) for biomarkers (also typically via elastic net) prior to training a second elastic net model on the phenotypic outcome or time-to-event ^14,17^. GrimAge is currently the gold-standard bAge epigenetic clock. It is derived from age, sex, and EpiScores of smoking pack years and 7 plasma proteins that have been associated with mortality or morbidity: adrenomedullin (ADM), beta-2-microglobulin (B2M), cystatin C, growth differentiation factor 15 (GDF15), leptin, plasminogen activation inhibitor 1 (PAI1), and tissue inhibitor metalloproteinase (TIMP1). Recently, a wider set of 109 EpiScores for the circulating proteome were generated by Gadd et al^14^. These have not yet been considered as potential features for the prediction of bAge.

Here, we improve the prediction of both cAge and bAge (**Figure 1**). We first present large-scale epigenome-wide association studies (EWAS) of age (for both linear and quadratic CpG effects) and all-cause mortality. A predictor of cAge is then generated using DNAm data from 13 cohorts, including samples from >18,000 participants of the Generation Scotland study^19^. We use a leave-one-cohort-out (LOCO) prediction framework, including dimensionality reduction prior to feature selection for linear and non-linear DNAm-age relationships (ascertained through the EWAS), and test it on ten external datasets. Through data linkage to death records, we develop a bAge predictor of all-cause mortality, which we compare against the current gold-standard predictor, GrimAge, in four external cohorts. These analyses highlight the potential for large DNAm resources to generate increasingly accurate predictors of (i) cAge, with potential forensic utility, and (ii) bAge, with potential implications for risk prediction and clinical trials.

**Figure 1.**
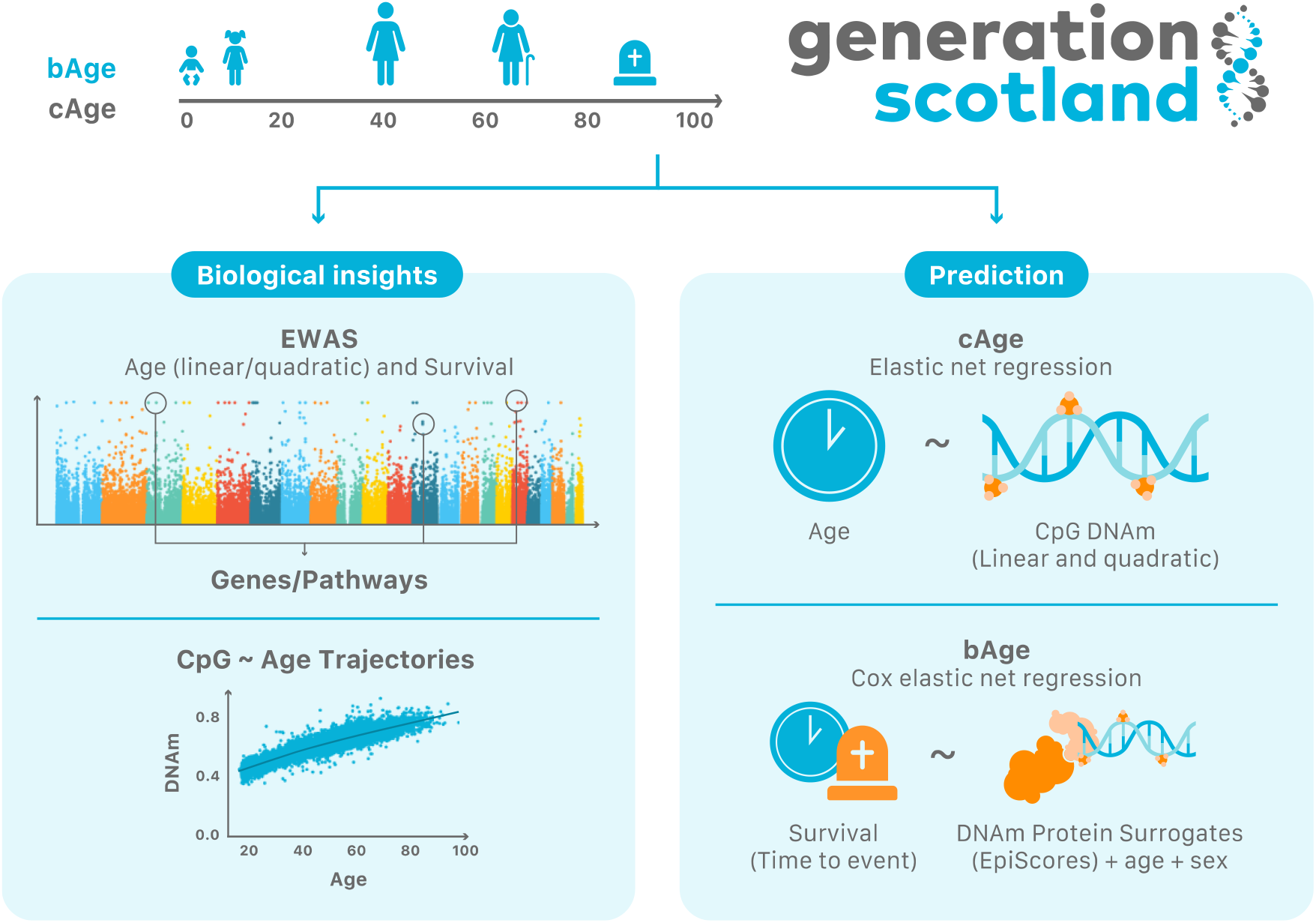
Study overview. Using the Generation Scotland cohort as our main data source, we explored the relationship between the epigenome and age/survival via EWAS, which also informed on genes of interest and potentially enriched pathways. We further characterised epigenome-wide CpG ~ age trajectories, which can be visualized in a new Shiny app, MethylBrowsR (https://shiny.igmm.ed.ac.uk/MethylBrowsR/). Finally, we refined epigenetic prediction of both cAge and bAge. Calculation of cAge can be performed either using a standalone script (https://github.com/elenabernabeu/cage_bage/tree/main/cage_predictor) or by uploading DNAm data to our MethylDetectR shiny app (https://shiny.igmm.ed.ac.uk/MethylDetectR/). As the weights for GrimAge and its component parts are not publicly available, bAge can only be calculated by using our standalone script (https://github.com/elenabernabeu/cage_bage/tree/main/bage_predictor), after obtaining GrimAge estimates from an external online calculator (http://dnamage.genetics.ucla.edu/new).

## Results

### Data overview

Generation Scotland is a Scottish family-based study with over 24,000 participants recruited between 2006 and 2011^19^. Blood-based DNAm levels at 752,722 CpG sites were quantified using the Illumina MethylationEPIC array for 18,413 individuals (see **Methods**). Participants were aged between 18 and 99 years at recruitment, with a mean age of 47.5 years (SD 14.9, **Table 1**). A total of 1,214 participant deaths have been recorded as of March 2022, via linkage to the National Health Service Central Register, provided by the National Records of Scotland.

**Table 1.** Age profile and test set prediction accuracy of cohorts used in cAge predictor training and testing. External cohort information taken from Zhang et al^5^. r column states Pearson correlation, RMSE the root mean squared error, and MAE the median absolute error.

| Cohort | N | Mean Age (SD) | Age Range | N<sub>Females</sub> (%) | Tissue | Prediction Accuracy |  |  |
| --- | --- | --- | --- | --- | --- | --- | --- | --- |
|  |  |  |  |  |  | r | RMSE | MAE |
| GS | 18,413 | 47.5 (14.9) | [17.1, 98.5] | 10,833 (58.8%) | Blood | - | - | - |
| LBC1921<sup>77,78</sup> | 692 | 82.3 (4.3) | [77.8,90.6] | 401 (57.9%) | Blood | 0.659 | 4.050 | 2.466 |
| LBC1936 | 2,795 | 73.6 (3.7) | [67.7,80.9] | 1,356 (48.5%) | Blood | 0.685 | 3.311 | 2.099 |
| GSE72775<sup>79</sup> | 335 | 70.2 (10.3) | [36.5, 90.5] | 138 (41.2%) | Blood | 0.949 | 3.275 | 1.843 |
| GSE78874<sup>79</sup> | 259 | 68.8(9.7) | [36.0, 88.0] | 113 (43.6%) | Saliva | 0.875 | 6.826 | 4.333 |
| GSE72773<sup>79</sup> | 310 | 65.6 (13.9) | [35.1, 91.9] | 150 (48.4%) | Blood | 0.945 | 4.611 | 2.068 |
| GSE72777<sup>79</sup> | 46 | 14.7 (10.4) | [2.2, 35.0] | 31 (67.4%) | Blood | 0.942 | 4.211 | 2.505 |
| GSE41169<sup>a,80</sup> | 95 | 31.6 (10.3) | [18.0, 65.0] | 28 (29.5%) | Blood | 0.975 | 2.869 | 1.947 |
| GSE40279<sup>4</sup> | 656 | 64.0 (14.7) | [19.0, 101.0] | 338 (51.5%) | Blood | 0.969 | 3.697 | 2.074 |
| GSE42861<sup>a,81</sup> | 689 | 51.9 (11.8) | [18.0, 70.0] | 492 (71.4%) | Blood | 0.972 | 4.498 | 3.563 |
| GSE53740<sup>a,82</sup> | 383 | 67.8(9.6) | [34.0, 93.0] | 155 (40.5%) | Blood | 0.921 | 4.443 | 2.797 |

<sup>a</sup> Some cohorts contain case/control data. GSE41169: Schizophrenia 62, control 33; GSE42861: Rheumatoid arthritis 354, control 335; GSE53740: Alzheimer's disease 15, corticobasal degeneration 1, frontotemporal dementia (FTD) 121, FTD/MND 7, progressive supranuclear palsy 43, control 193, unknown 4.

In order to train and test a cAge predictor, data from an additional 6,260 individuals from ten external cohorts were considered. These included the Lothian Birth Cohorts (LBC) of 1921 and 1936, and eight publicly available Gene Expression Omnibus (GEO) datasets (see Methods, **Table 1**). Given that the external datasets assessed DNAm (blood-based apart from GSE78874, which considered saliva) using the Illumina HumanMethylation450K array, the Generation Scotland data were subset to 374,791 CpGs that were present across all studies.

To test the bAge predictor, data from an additional 4,134 individuals (with a total of 1,653 deaths) from six external cohorts were considered. These included both the LBC1921 and LBC1936 cohorts, as well as the Framingham Heart Study (FHS) and the Women’s Health Initiative (WHI) Broad Agency Award 23 (B23) study for Black, White, and Hispanic individuals (see Methods, **Table 2**).

**Table 2.**
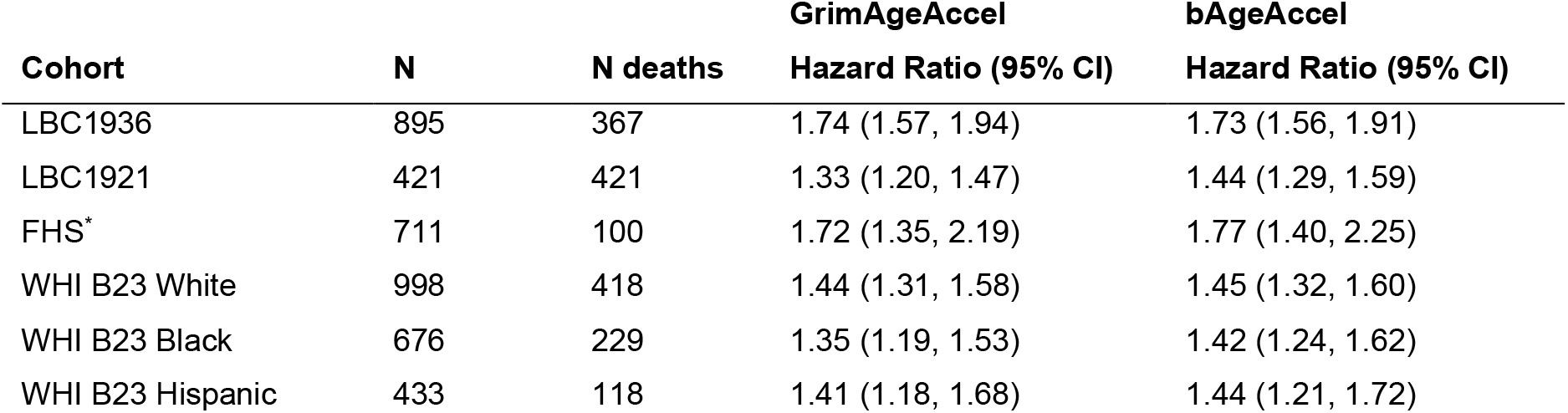
Cox Proportional Hazards output for GrimAgeAccel and bAgeAccel in the test datasets. Hazard ratios are presented per standard deviation of the GrimAgeAccel and bAgeAccel variables. Further details in Supplementary Table 11. *The FHS cohort used here was the same as the test set from the original GrimAge paper.

| Cohort | N | N deaths | GrimAgeAccel | bAgeAccel |
| --- | --- | --- | --- | --- |
|  |  |  | Hazard Ratio (95% CI) | Hazard Ratio (95% CI) |
| LBC1936 | 895 | 367 | 1.74 (1.57, 1.94) | 1.73 (1.56, 1.91) |
| LBC1921 | 421 | 421 | 1.33 (1.20, 1.47) | 1.44 (1.29, 1.59) |
| FHS* | 711 | 100 | 1.72 (1.35, 2.19) | 1.77 (1.40, 2.25) |
| WHI B23 White | 998 | 418 | 1.44 (1.31, 1.58) | 1.45 (1.32, 1.60) |
| WHI B23 Black | 676 | 229 | 1.35 (1.19, 1.53) | 1.42 (1.24, 1.62) |
| WHI B23 Hispanic | 433 | 118 | 1.41 (1.18, 1.68) | 1.44 (1.21, 1.72) |

### Epigenome-wide association studies of cAge

EWAS of cAge were performed in the Generation Scotland cohort, resulting in 99,832 linear and 137,195 quadratic CpG associations that were epigenome-wide significant (*p* < 3.6 × 10^−8^, **Supplementary Figure 1, Supplementary Table 1 and 2**, see **Methods**). These mapped to 17,339 and 19,432 unique genes, respectively. There were 48,312 CpGs with both a significant linear and quadratic association.

The most significant linear associations included cg16867657 and cg24724428 (*ELOVL2*), cg08097417 (*KLF14*), and cg12841266 (*LHFPL4)*, all *p* < 1 × 10^−300^, (**Supplementary Table 1, Supplementary Figure 2**). Around half of the CpGs with a significant linear association (51,213/99,832, 51.3%) showed a positive association with age. The most significant quadratic associations were cg11084334 (*LHFPL4, p* = 6.49 × 10^−206^), cg15996534 (*LOC134466, p* = 8.7 × 10^−194^), and cg23527621 (*ECE2* and *CAMK2N2, p* = 9.95 × 10^−189^, **Supplementary Table 2, Supplementary Figure 3**).

The univariate associations between all 752,722 CpGs and cAge in a subset of 4,450 unrelated participants (DNAm arrays processed together in a single experiment) from Generation Scotland can be visualised via an online ShinyApp, MethylBrowsR (https://shiny.igmm.ed.ac.uk/MethylBrowsR/).

### Prediction of cAge

Epigenetic clocks for cAge were created using elastic net penalised regression. Input features consisted of CpG and CpG^2^ DNAm values for sites that were epigenome-wide significant in their corresponding EWAS analysis (see **Methods, Figure 2**). After iterating through combinations of CpG and CpG^2^ terms (ranked by EWAS p-value), the best-performing model considered the top 10,000 CpG and top 300 CpG^2^ sites from the EWAS as potentially informative features (see **Methods, Supplementary Table 3 and 4, Supplementary Figure 4 and 5**). A single external cohort was used for this screening step (GSE40279, N = 656) and model fit was based on the root mean squared error (RMSE) and median absolute error (MAE) of prediction.

**Figure 2.**
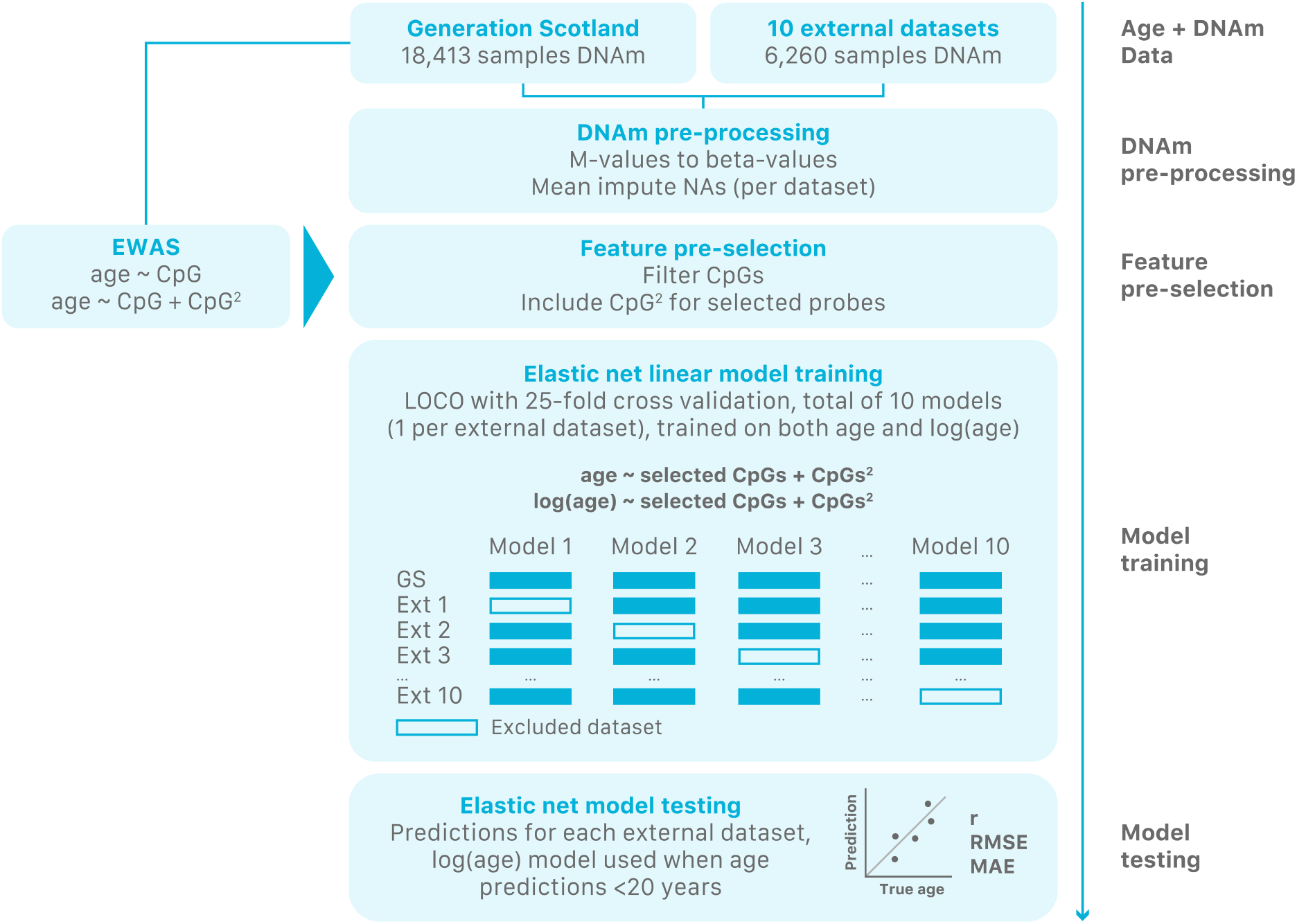
Flowchart for the creation of the cAge predictor. First, DNAm data originating from Generation Scotland and 10 external datasets was pre-processed. Next, CpGs were pre-selected based on the Generation Scotland EWAS for genome-wide significant linear and quadratic CpG-age associations. Elastic net models were then trained and tested on the remaining features using a LOCO framework with 25-fold cross validation, with training on both age and log(age) as outcomes.

A LOCO framework was used to train the cAge predictor, whereby for each of the 10 external cohorts, a model was trained on data from Generation Scotland and the remaining nine external cohorts. Testing was then performed on the excluded cohort (total N_testing_ = 6,260). A final model was also trained on all 11 cohorts (N_training_ = 24,673).

Both age and log(age) were considered as outcomes, with the latter showing better prediction results in younger individuals, reflecting the importance of considering non-linear DNAm-age associations in cAge prediction. As a result, if the initial cAge prediction was <20 years, that individual’s predicted age was re-estimated using weights from the log(age) model.

The combined LOCO prediction results showed a strong correlation with cAge (r = 0.96, **Figure 3, Table 1**) and a MAE of 2.3 years. Furthermore, 24% of individuals were classified to within one year of their chronological age. The cohort with the largest prediction errors was GSE78874, in which DNAm was measured in saliva instead of blood.

**Figure 3.**
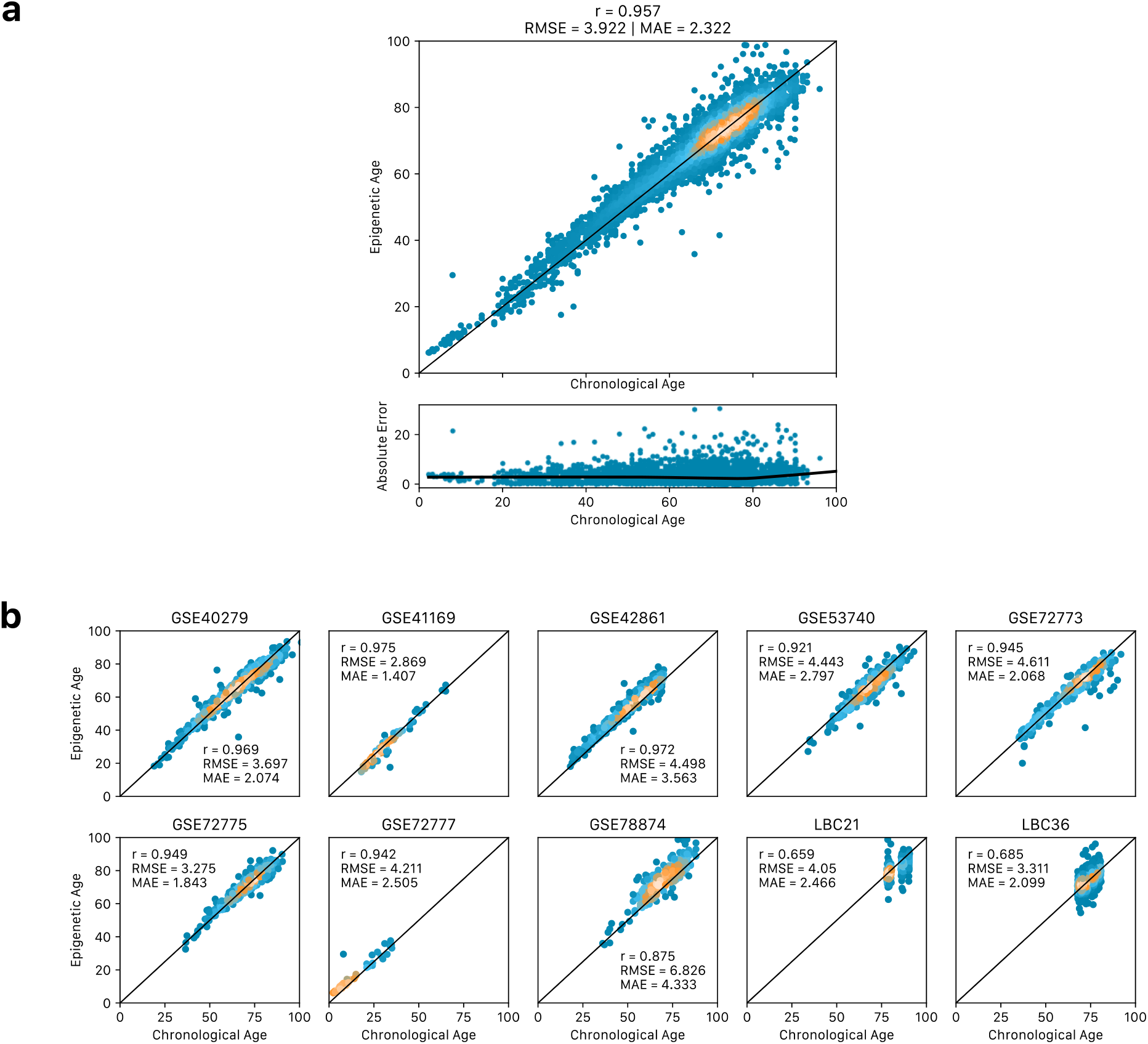
cAge predictor performance on 10 external testing datasets, (a) across all datasets considered, and (b) per cohort. Performance metrics shown include Pearson correlation (r), root mean squared error (RMSE), and median absolute error (MAE). Metrics also included in **Table 1**.

The elastic net model (trained in all 11 cohorts) with the lowest mean cross-validated error identified 2,330 features (2,274 linear and 56 quadratic) as most predictive of age, and 1,986 features (1,931 linear and 55 quadratic) as most predictive of log(age). The weights for the age model are presented in **Supplementary Table 5**, and for the log(age) model in **Supplementary Table 6**.

### Epigenome-wide association study of all-cause mortality

To identify individual CpG loci associated with survival, an EWAS on time to all-cause mortality was performed in Generation Scotland (N_deaths_ = 1,214, see **Methods**). This analysis identified 1,182 epigenome-wide significant associations (*p* < 3.6 × 10^−8^, **Supplementary Figure 6**), which mapped to 704 unique genes. Around a third (418/1,182 = 35.36%) of these CpGs were associated with a decreased survival time. The lead findings included CpGs mapping to smoking-related loci^10,20–24^ such as cg05575921 (*AHRR, p* = 3.01 × 10^−57^), cg03636183 (*F2RL3, p* = 6.78 × 10^−44^), cg19859270 (*GPR15, p* = 1.09 × 10^−33^), cg17739917 (*RARA, p* = 1.92 × 10^−33^), cg14391737 (*PRSS23, p* = 5.59 × 10^−33^), cg09935388 (*GFI1, p =* 3.30 × 10^−31^), and cg25845814 (*ELMSAN1/MIR4505, p* = 1.31 × 10^−30^) (**Supplementary Table 7**). Of the non-smoking-related CpGs amongst the top 50 associations, seven mapped to genes whose methylation has been linked to various forms of cancer, including *ZMIZ1*^25^, *SOCS3*^26–28^, *ZMYND8*^29^ and *CHD5*^30–32^. Another probe mapped to *FKBP5*, a gene whose methylation is involved in the regulation of the stress response, and which has been linked to increased cardiometabolic risk through accelerated ageing^33^. Finally, one top probe mapped to *SKI*, whose methylation has been linked to age-related macular degeneration^34^. All associations remained after adjusting for relatedness in the Generation Scotland cohort (see **Methods, Supplementary Table 8**).

There was a high correlation of the Z-score effect sizes across the 200 sites that overlapped between our study and the 257 epigenome-wide significant findings from a recent large (N = 12,300, N_deaths_ = 2,561) meta-analysis of all-cause mortality (r = 0.58, **Supplementary Figure 7**). All 200 sites were significant at a nominal *p* < 0.05 threshold and 25 were epigenome-wide significant at *p* < 3.6 × 10^−8^.

A gene-set enrichment analysis considering genes to which epigenome-wide significant CpGs mapped to returned 198 significantly enriched (FDR *p* < 0.05) GO biological processes (see **Methods**, full FUMA gene-set enrichment results in **Supplementary Table 9**). The most significantly enriched GO terms included processes relating to neurogenesis/neuron differentiation and development, positive immune system regulation and development, cell motility and organization, and regulation of protein modification/phosphorylation. Other significantly enriched sets included sites bound by FOXP3, ETS2, and the PML-RARA fusion protein.

### Prediction of bAge

Amongst the second generation epigenetic clocks, GrimAge is the current best predictor of lifespan (time to death)^17^. In an effort to improve the prediction of bAge, an elastic net Cox model was trained on all-cause mortality in Generation Scotland (N_total_ = 18,365, N_deaths_ = 1,214, see **Methods**). The GrimAge components (age, sex, and EpiScores for smoking and 7 plasma proteins) and Gadd et al’s 109 protein EpiScores^14^ were considered as potentially-informative features (**Figure 4**).

**Figure 4.**
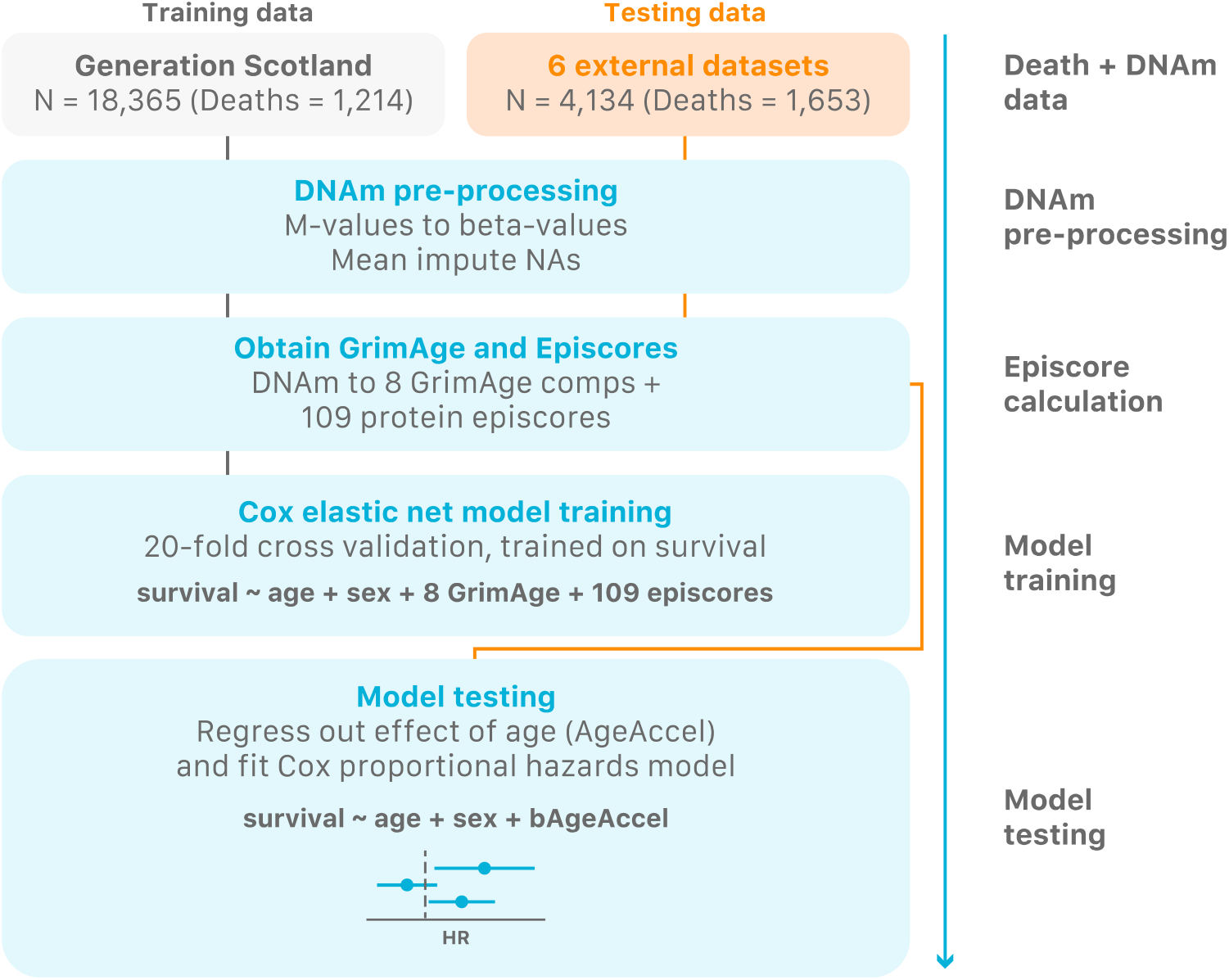
Flowchart for the creation of the bAge predictor. First, DNAm data originating from Generation Scotland and six external datasets was pre-processed. GrimAge components and 109 protein EpiScores were generated within each cohort. A Cox proportional hazards elastic net regression model of all-cause mortality (with 20-fold cross validation) was trained in Generation Scotland with the GrimAge components and EpiScores as possible features. The model that maximised Harrell’s C index was tested on the six external datasets.

The elastic net Cox model identified a weighted sum of 35 features as most predictive of all-cause mortality in Generation Scotland. These included age and the GrimAge smoking EpiScore, along with 5/7 protein EpiScores from GrimAge (B2M, cystatin C, GDF15, PAI1, and TIMP1), and 28/109 protein EpiScores from Gadd et al^14^. Amongst these were EpiScores for C-reactive protein (CRP), the growth hormone receptor (GHR) protein, and numerous cytokines (CCL11, CCL23, CCL18, CXCL10, CXCL9, CXCL11, and HGF). The weights for the linear predictor are presented in **Supplementary Table 10**.

The bAge predictor was regressed on age to obtain a measure of epigenetic age acceleration (bAgeAccel). The epigenetic age acceleration residuals showed significant associations with all-cause mortality across four test cohorts of differing ancestries (**Table 2, Supplementary Table 11, Figure 5**). The bAge measure showed slightly stronger associations than GrimAge (also regressed on age, termed GrimAgeAccel) in fixed effects meta-analyses (Hazard Ratio and 95% Confidence Interval per SD difference of GrimAgeAccel and bAgeAccel: HR = 1.47 [1.40, 1.54] with *p* = 1.08 × 10^−52^, and HR = 1.52 [1.44, 1.59] with *p* = 2.20 × 10^−60^, respectively.

**Figure 5.**
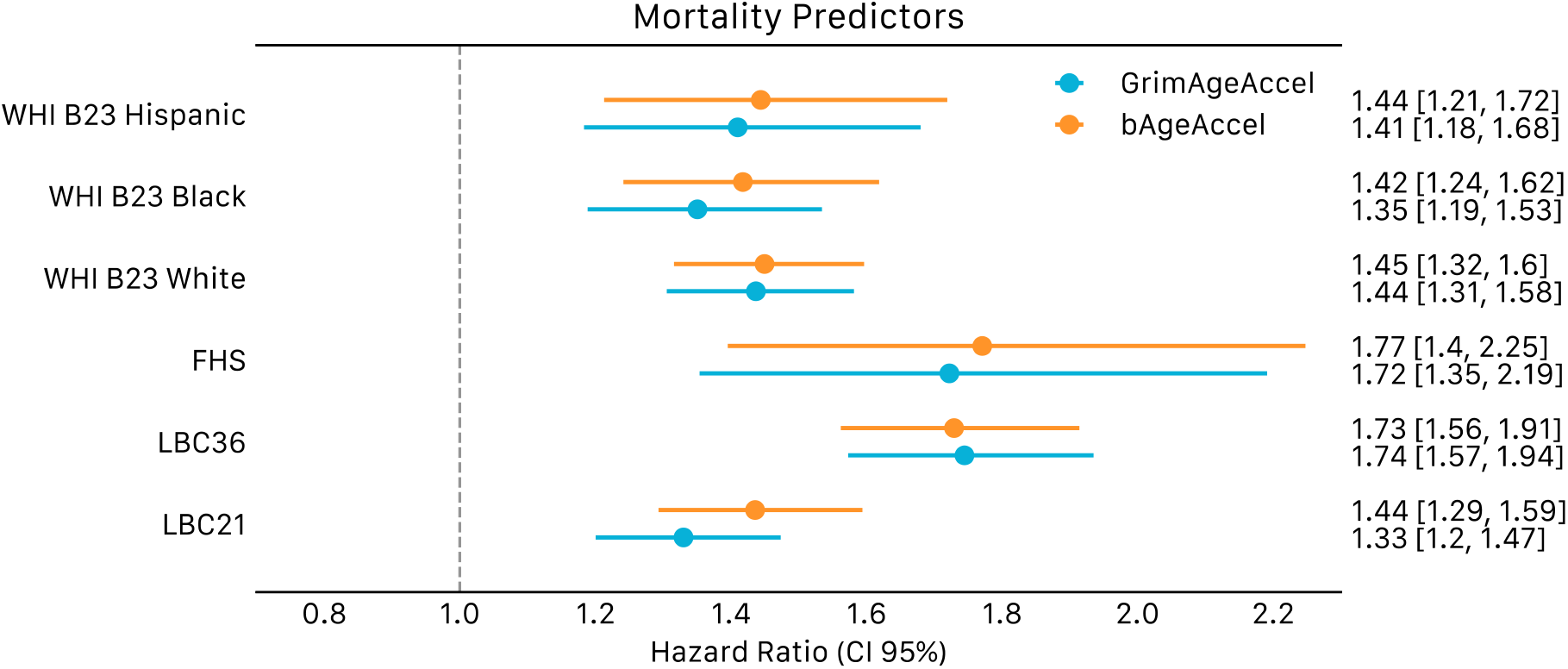
Forest plots of bAge/GrimAge predictors, applied to all-cause mortality in LBC1921, LBC1936, FHS, and WHI. Predictors regressed on age. Hazard ratios are presented per standard deviation of the GrimAgeAccel and bAgeAccel variables, along with 95% confidence intervals. Cox models are adjusted for age at DNAm sampling and sex.

## Discussion

Accurate predictors of cAge and bAge have major implications for biomedical science and healthcare through risk prediction and preventative medicine. Here, we present improved DNAm-based predictors of age and lifespan.

Epigenetic cAge prediction is expected to reach near-perfect estimates as sample sizes grow^5^. Making use of Generation Scotland, a very large single-cohort DNAm resource, we derived a cAge predictor with a MAE of 2.3 years, tested in over 6,000 external samples. Our predictor has potential forensic applications, although ethical caveats exist^8^. In addition, despite the high correlations and low RMSE and MAE estimates at the population level, there are still several individuals with inaccurate predictions (e.g. > 20 years between predicted and actual age, **Figure 3**), though this could also reflect sample mix-ups or data entry errors.

cAge prediction was improved when accounting for non-linear relationships between DNAm and age. Whilst generally understudied, non-linear patterns have been found at numerous CpG sites, where DNAm is found to increase rapidly in early ages and stabilize in adulthood, potentially reflecting developmental processes^35^. Similarly, stable DNAm levels followed by rapid methylation/demethylation have also been described in later life^36^, which could offer insight into aging-specific processes. Given the number of samples from individuals aged 20 or under in the training of our predictor (N=574/24,674= 2.4%), we may not have captured the full extent of DNAm-based ageing patterns in the younger population. Future studies could also consider sex-specific models as diverging non-linear patterns between males and females have been shown in previous studies^37^. Interactions between CpGs along with higher order polynomial terms and spline-based models might better capture some of these non-linear changes.

The development of the cAge predictor highlighted the advantages of feature pre-selection ahead of penalised elastic net regression. Compared to a model with all possible features in the training set (r = 0.93, RMSE = 5.25, MAE = 3.43, pre-selection greatly improved performance (r = 0.96, RMSE = 3.92, MAE = 2.32). Several DNAm studies of age and age-related phenotypes have used pre-selection methods (e.g., filtering by magnitude of correlation or strength of association) instead of, or in addition to elastic net^38–45^. Whereas the feature pre-selection here required arbitrary decisions on thresholds, other studies have found that feature reduction via PCA optimises DNAm predictors^46,47^.

Feature pre-selection may have aided cAge predictions by screening out CpGs with low intra-sample variability due to technical variance^48,49^. One previous study^47^ observed that CpGs with stronger cAge associations were more reliable. A limitation of our approach to feature pre-selection was that it was biased towards the Generation Scotland cohort in which the age EWAS were conducted. We also note that pre-selection introduces statistical challenges associated to post-selection inference^50^. Furthermore, our penalised regression modelling strategy for cAge only incorporated additive effects. Non-additive tree ensemble methods and other machine learning frameworks may improve predictions further^51^. Finally, as our predictor has been mainly trained and tested on blood data, it may not generalise to other tissues.

Whilst a single DNAm predictor of cAge is of interest, the selected CpG features are unlikely to identify all epigenome-wide patterns related to ageing. Our EWAS of chronological age identified 99,832 linear and 137,195 quadratic CpG-age associations. The sample size was more than double that of the largest study reported on the EWAS Catalog^52^ - our previous Generation Scotland analysis^53^. In addition to refining our previously described DNAm-age linear associations, we have extended previous small-scale approaches to highlight non-linear patterns^36,37^. As shown here, these findings can aid the predictive performance of epigenetic clocks, and may additionally improve our understanding of epigenetic changes during development and ageing-related decline in later life.

Recent work has shifted focus from the prediction of cAge to bAge, with more expansive clinical applications. Our new bAge predictor of all-cause mortality had a greater effect size and was more statistically significant than GrimAge in the external test set meta-analysis. GrimAge is already being used as an end-point for clinical trials^54^ and studies of rejuvenation^55,56^. The bAge predictor included EpiScores for CRP and numerous cytokines, which reflect inflammation and predict overall and cardiovascular mortality^57–59^. Chronic inflammation can lead to several diseases, including cardiovascular disease and exacerbates the ageing process^60,61^. In addition, the growth hormone receptor (GHR) protein EpiScore was selected; both the receptor and its corresponding protein have been linked to longevity in mouse models^62–66^. 25/28 of the selected EpiScores from Gadd et al^14^ have been associated to multiple diseases, including diabetes, chronic obstructive pulmonary disease, ischaemic heart disease, lung cancer, Alzheimer’s, rheumatoid arthritis, stroke, and depression (**Supplementary Table 10**). As sample sizes for cause-specific mortality outcomes increase, a more granular suite of lifespan predictors can be developed.

Whereas the cAge predictions translated into external cohorts with minimal calibration issues, individual-level bAge predictions were highly variable. Future work for these (and all) DNAm array-based predictors should consider the limitations of signatures that lack absolute thresholds/cut-points for risk prediction in a new individual selected at random from the population.

A total of 1,182 epigenome-wide significant associations were identified in our EWAS of all-cause mortality. The most significant probes mapped to genes previously associated with smoking, such as *AHRR, F2RL3*, and *GPR15*^67^. Hypomethylation at probes nearby these genes has been previously linked to increased mortality risk, be that all-cause or disease specific (e.g., cancer or, cardiovascular-related mortality)^20,68–70^. Other, non-smoking related, lead probes mapped to genes whose methylation has been linked to various forms of cancer, increased cardiometabolic risk, and age-related macular degeneration^25–34^. There was moderate agreement (correlation of 0.58 between Z scores) between our findings and the significant results from a previous EWAS meta-analysis of survival. However, different covariates and ancestries were considered across these studies. An enrichment analysis highlighted links to neurodevelopment and immune regulation, as well as to sites bound by FOXP3, ETS2, and the PML-RARA fusion protein. FOXP3 is a transcriptional regulator involved in the development and inhibitory function of regulatory T-cells^71^. ETS2 and PML-RARA are a protooncogene and a protein resulting from a chromosomal translocation that resulting in an oncofusion protein, respectively, having both been linked to acute myeloid leukemia^72,73^. This finding may be influenced by the large number of cancer-related deaths in Generation Scotland (N = 509). Further work is needed to disentangle the role of methylation/demethylation at these sites with survival. Future EWAS on specific mortality causes will highlight mechanisms underlying age- and disease-related decline.

The integration of multiple large datasets and new approaches to feature selection has facilitated improvements to the blood-based epigenetic prediction of biological and chronological age. The inclusion of multiple protein EpiScore features and consideration of quadratic DNAm effects may also be relevant for other EWAS and prediction studies. Together, this can improve our biological understanding of complex traits and the prediction of adverse health outcomes.

## Methods

### Generation Scotland

#### Cohort description

Generation Scotland: Scottish Family Health Study is a population-based cohort study that includes ~7,000 families from across Scotland^19^. Study recruitment took place between 2006 and 2011 when participants were aged between 17 and 99 years (**Table 1**). In addition to completing health and lifestyle questionnaires, participants donated blood or saliva samples for biomarker and omics analyses. The majority of participants also provided consent for linkage to their electronic medical records, yielding retrospective and prospective information on primary and secondary disease outcomes as well as prescription data.

#### Data linkage to death records

Information on mortality and cause of death is routinely updated via linkage to the National Health Service Central Register, provided by the National Records of Scotland. The data used here were correct as of March 2022, with a total of 1,214 deaths and 18,365/18,413 samples with non-missing and non-negative time-to-death/event (TTE) values. Average TTE amongst deaths was 7.79 (SD 3.54) years. Leading causes of death included malignant neoplasms (509), ischaemic heart disease (134), cerebrovascular disease (69), other forms of heart disease (44), chronic lower respiratory disease (42), mental disorders including dementia (36), and other degenerative diseases of the nervous system (35).

#### DNA methylation in Generation Scotland

DNA methylation in blood was quantified for 18,413 Generation Scotland participants across three separate sets (N_Set1_ = 5,087, N_Set2_ = 4,450, N_Set3_ = 8,876) using the Illumina MethylationEPIC (850K) array. Individuals in Set 1 included a mixture of related and unrelated individuals. Set 2 comprised individuals unrelated to each other and also to those in Set 1. Set 3 contained a mix of related individuals – both to each other and to those in Sets 1 and 2 – and included all remaining samples available for analysis.

Quality control details have been reported previously^53,74^. Briefly, probes were removed based on (i) outliers from visual inspection of the log median intensity of the methylated versus unmethylated signal per array, (ii) a bead count < 3 in more than 5% of samples, (iii) ≥ 5% of samples having a detection *p*-value > 0.05, (iv) if they pertained to the sex chromosomes, (v) if they overlapped with SNPs, and/or (vi) if present in potential cross-hybridizing locations^75^. Samples were removed (i) if there was a mismatch between their predicted sex and recorded sex, (ii) if ≥ 1% of CpGs had a detection *p*-value > 0.05, (iii) if sample was not blood-based, and/or (iv) if participant responded “yes” to all self-reported diseases in questionnaires. Dasen normalisation^76^ was carried out per set (for cAge training) or across all individuals (for EWAS). A total of 752,722 CpGs remained after QC. To maximise the generalisability of the predictors across different versions of Illumina arrays, we subset the content to the intersection of sites on the EPIC and 450k arrays, as well as to those present across all cohorts considered in the study (**Table 1**), totalling 374,791 CpGs.

### External datasets

To test the cAge predictor, we considered DNA methylation for a total of 6,260 external samples, from eight publicly available datasets from the Gene Expression Omnibus (GEO) resource and repeated measures (up to four time points) from two cohorts of blood-based DNAm, the Lothian Birth Cohorts (LBC) of 1936 and 1921 (**Table 1**)^4,77–82^. The baseline samples from the LBC cohorts, along with the Framingham Heart Study (FHS) and the Women’s Health Initiative (WHI) study, were also used for the testing of our bAge predictor (**Table 2**).

#### Lothian Birth Cohorts

LBC1921 and LBC1936 are longitudinal studies of ageing on individuals born in 1921 and 1936, respectively^77^. Study participants completed the Scottish Mental Surveys of 1932 and 1947 at approximately age 11 years old and were living in the Lothian area of Scotland at the time of recruitment in later life. Blood samples considered here were collected at around age 79 for LBC1921, and at around age 70 for LBC1936. DNA methylation was quantified using the Illumina HumanMethylation450 array, for a total of 692 (up to 3 repeated measurements from 469 individuals) and 2,795 (up to 4 repeated measurements from 1,043 individuals) samples from LBC1921 and LBC1936 respectively. Quality control details have been reported previously^5,83^. Briefly, probes were removed (i) if they presented a low (< 95%) detection rate with *p-*value < 0.01, and/or (ii) if they presented inadequate hybridization, bisulfite conversion, nucleotide extension, or staining signal, as assessed by manual inspection. Samples were removed (i) if they presented a low call rate (<450,000 probes detected at *p*-value < 0.01) and/or (ii) if predicted sex did not match reported sex. Finally, as stated previously, probes were filtered down to the 374,791 common across all datasets (**Table 1**). Missing values were mean imputed.

A total of 421 and 895 samples from LBC1921 and LBC1936 respectively, corresponding to the first wave of each study (thus aged around 79 and 70 at time of sampling for each cohort respectively), were used in our bAge analysis (**Table 2**). All-cause mortality was assessed via linkage to the National Health Service Central Register, provided by the National Records of Scotland. The data used here are correct as of January, 2022, with a total of 421 and 367 deaths in LBC1921 and LBC1936 respectively.

#### Gene Expression Omnibus (GEO) datasets

DNAm and age information for 2,773 individuals from a total of 8 datasets was downloaded from the public domain (Gene Expression Omnibus, GEO). DNAm was quantified with Illumina’s HumanMethylation450 chip. QC information can be found in each pertaining publication (**Table 1**), and CpGs were filtered down to the 374,791 common across all datasets. Missing values were mean imputed.

#### Framingham Heart Study (FHS)

The FHS cohort is a large-scale longitudinal study started in 1948, initially investigating the common factors of characteristics that contribute to cardiovascular disease (CVD)^84^. The study at first enrolled participants living in the town of Framingham, Massachusetts, who were free of overt symptoms of CVD, heart attack or stroke at enrolment. In 1971, the study established the FHS Offspring Cohort to enrol a second generation of the original participants’ adult children and their spouses for conducting similar examinations^85^. Participants from the FHS Offspring Cohort were eligible for our study if they attended both the seventh and eighth examination cycles and consented to having their molecular data used for study. We used data pertaining to a total of 711 individuals which had not been used in the training of GrimAge, and for which DNAm data and death records were available. Peripheral blood samples were obtained on the eight examination cycle, and DNAm data was measured using the Illumina Infinium HumanMethylation450 array, with QC details are described elsewhere^17^. Deaths recorded are accurate as of 1st January 2013, with a total of 100 recorded.

#### Women’s Health Initiative (WHI)

The WHI study enrolled postmenopausal women aged 50-79 years into the clinical trials (CT) or observational study (OS) cohorts between 1993 and 1998. We included 2,107 women from “Broad Agency Award 23” (WHI BA23). WHI BA23 focuses on identifying miRNA and genomic biomarkers of coronary heart disease (CHD), integrating the biomarkers into diagnostic and prognostic predictors of CHD and other related phenotypes. This cohort is divided into three datasets, pertaining to three different ancestries: White, Black, and Hispanic, with 998, 676, and 433 participants respectively. Blood-derived DNAm data was available for participants. DNAm data was measured using the Illumina Infinium HumanMethylation450 array, QC details described elsewhere^17^. Deaths recorded are accurate as March 1^st^, 2017, with a total of 418, 229, and 118 recorded for White, Black, and Hispanic ancestries respectively.

### EWAS of chronological age

We conducted an EWAS to identify CpG sites that had linear or quadratic associations with chronological age, using Generation Scotland data (N = 18,413, CpGs = 752,722). Linear regression analyses were carried out which included both linear and quadratic CpG M-values as predictor variables and age as the dependent variable (Age ~ CpG and Age ~ CpG + CpG^2^, respectively). Fixed effect covariates included estimated white blood cell (WBC) proportions (basophils, eosinophils, natural killer cells, monocytes, CD4T, and CD8T cells) calculated in the *minfi* R package (version 1.36.0)^86^ using the Houseman method^87^, sex, DNAm batch/set, smoking status (current, gave up in the last year, gave up more than a year ago, never, or unknown), smoking pack years, and 20 methylation based principal components (PCs) to correct for unmeasured confounders. Age was centered by its mean, and CpG and CpG^2^ M-values were scaled to mean zero and variance one. Epigenome-wide significance was set at *p*-value < 3.6 × 10^−8^, as per Saffari et al^88^.

### Prediction of chronological age

Elastic net regression (with α = 0.5 as the L_1_, L_2_ mixing parameter) was used to derive a predictor of chronological age from the 374,791 CpG sites common across all cohorts considered in cAge training (description of cohorts in **Table 1**). The *biglasso* R package (version 1.5.1) was used^89^, with 25-fold cross validation (CV) to select the shrinkage parameter (λ) that minimised the mean cross-validated error. This resulted in randomly assigned folds of ~1,000 individuals. A sensitivity analysis was performed, assigning individuals from the same methylation set and cohort to individual folds, which returned highly similar results.

#### Leave-one-cohort-out (LOCO)

The cAge predictor was created and tested using a leave-one-cohort-out (LOCO) framework, where the model was trained in 10 cohorts and tested on the excluded external cohort (**Figure 2**). The final reported model was trained using all 11 sets described here. Pearson correlations (r) with reported age were calculated along with the root mean square error (RMSE) and median absolute error (MAE).

#### Log(age)

In addition to training on chronological age, models were also trained on the natural logarithm of chronological age, log(age). The age of our test samples was predicted using the model fit on chronological age, and, if the predicted age returned was 20 years or younger, a new prediction was obtained making use of the model fit on log(age). This approach parallels that in Horvath’s 2013 clock, which log-transforms chronological age in under 20s prior to training^3^.

#### Feature pre-selection

Several studies have highlighted the benefits of feature pre-selection for elastic net^46,47^. Here, we performed preliminary analyses, including differently sized subsets of CpG sites as features in elastic net. We considered sites that were epigenome-wide significant at *p* < 3.6 × 10^−8^ and then ranked CpGs in ascending order of *p*-value (most significant ranked first), before defining subsets of varying sizes (from 1,000 to 300,000 CpGs). Our training cohort was Generation Scotland, whilst our test set was GSE40279, one of the largest external datasets with the widest age range. Our analyses showed that the 10,000 most significant loci (age - CpG associations) yielded the test set predictions with the highest r and lowest RMSE (**Supplementary Table 3, Supplementary Figure 4**). In addition to these sites, subsets of CpGs with a significant quadratic relationship to age were explored, with subset sizes varying from 100 to 20,000. These features were included in training as CpG^2^ beta values, and, when not already present in the model, in their linear form as well. In addition to the top 10,000 age-associated CpGs, the top 300 quadratic sites from our EWAS yielded the best performing model (**Supplementary Table 4, Supplementary Figure 5**). This final list of features was then trained and tested using a LOCO framework, as described above.

While this involves substantial overfitting in the training data, the test sets (other than GSE40279) remained completely independent prior to the prediction analyses.

### EWAS of all-cause mortality

An EWAS was conducted to identify CpG sites (from a total of 752,722 loci) that were associated with time to all-cause mortality in Generation Scotland. Cox Proportional Hazards (Cox PH) regression models were fit for each CpG site as predictor of interest using the *coxph* function from the *survival* R package (version 3.3.1), with time-to-death or censoring as the survival outcome. Fixed effect covariates included those used in our cAge EWAS (age at baseline, sex, set/batch, smoking status, smoking pack years, WBC estimates, and top 20 methylation PCs). Epigenome-wide significance was set at *p*-value < 3.6 × 10^−8^.

To assess whether relatedness in the cohort influenced the results, a Cox PH model with a kinship matrix was fit for each significantly associated CpG, using the *coxme* R package (version 2.2.16). All associations were replicated at *p* < 3.6 × 10^−8^ (**Supplementary Table 8**).

### Prediction of survival (biological age)

#### Training in Generation Scotland

To train a bAge predictor, component scores for GrimAge were estimated for all Generation Scotland samples via Horvath’s online calculator^17^ (http://dnamage.genetics.ucla.edu/new). These included DNAm estimates of smoking and seven proteins – DNAm ADM, DNAm B2M, DNAm cystatin C, DNAm GDF15, DNAm leptin, DNAm PAI1, and DNAm TIMP1. Each variable was then standardised to have a mean of zero and variance of one. We also considered DNAm EpiScores for 109 proteins as described by Gadd et al^14^. The 109 EpiScores were projected into Generation Scotland via the MethylDetectR^90^ Shiny App (https://shiny.igmm.ed.ac.uk/MethylDetectR/) before being standardised to have a mean of zero and variance of one.

This resulted in 116 protein EpiScores, a smoking EpiScore, plus chronological age and sex as features for an elastic net Cox PH model (R package *glmnet* version 4.1.4). 20-fold CV was performed (with approximately 1,000 individuals per fold), with individuals from the same batch/set included in the same fold, and with Harrell’s C index used to evaluate the optimal λ value.

#### Testing in LBC, FHS, and WHI

The association between bAgeAccel (the residual of bAge regressed on chronological age to obtain measure of accelerated epigenetic ageing) and mortality was assessed in six datasets from four external studies: LBC1921 and LBC1936, FHS, and the WHI studies for White, Black, and Hispanic ancestries (**Table 2**). After generating the bAge predictors in the external datasets, Cox proportional hazards models, adjusting for age and sex, were used to compare associations with all-cause mortality for GrimAgeAccel and bAgeAccel. We examined Schoenfeld residuals in the LBC models to check the proportional hazards assumption at both global and variable-specific levels using the *cox*.*zph* function from the R *survival* package (version 3.3.1). We restricted the TTE period by each year of possible follow-up, from 5 to 21 years, and found minimal differences in the bAgeAccel-survival HRs between follow-up periods that did not violate the assumption and those that did (**Supplementary Table 12**).

### Enrichment analyses

A gene set enrichment analysis was performed using the Functional Mapping and Annotation (FUMA) GENE2FUNC tool^91^, which employs a hypergeometric test. Background genes employed included all unique genes tagged by CpGs in the EPIC array. FDR *p*-value threshold was set at 0.05, and the minimum number of overlapping genes within gene sets was set to 2.

## Ethics

All components of Generation Scotland received ethical approval from the NHS Tayside Committee on Medical Research Ethics (REC Reference Number: 05/S1401/89). Generation Scotland has also been granted Research Tissue Bank status by the East of Scotland Research Ethics Service (REC Reference Number: 20-ES-0021), providing generic ethical approval for a wide range of uses within medical research.

Ethical approval for the LBC1921 and LBC1936 studies was obtained from the Multi-Centre Research Ethics Committee for Scotland (MREC/01/0/56) and the Lothian Research Ethics committee (LREC/1998/4/183; LREC/2003/2/29). In both studies, all participants provided written informed consent. These studies were performed in accordance with the Helsinki declaration.

## Availability of data and material

According to the terms of consent for Generation Scotland participants, access to data must be reviewed by the Generation Scotland Access Committee. Applications should be made to.

Lothian Birth Cohort data are available on request from the Lothian Birth Cohort Study, University of Edinburgh (https://www.ed.ac.uk/lothian-birth-cohorts/data-access-collaboration). Lothian Birth Cohort data are not publicly available due to them containing information that could compromise participant consent and confidentiality.

All custom R (version 4.0.3), Python (version 3.9.7), and bash code is available with open access at the following GitHub repository: https://github.com/elenabernabeu/cage_bage

EWAS summary statistics will be submitted to the EWAS catalog upon acceptance. They are currently available for open access on Edinburgh DataShare: https://datashare.ed.ac.uk/handle/10283/4496 cAge predictions can be obtained using MethylDetectR (https://shiny.igmm.ed.ac.uk/MethylDetectR/) or via a standalone script: https://github.com/elenabernabeu/cage_bage/tree/main/cage_predictor

As the CpG weights for the GrimAge components are not publicly available, bAge predictions first require users to generate GrimAge estimates from the following online calculator (http://dnamage.genetics.ucla.edu/new). bAge can then be estimated via the following standalone script: https://github.com/elenabernabeu/cage_bage/tree/main/bage_predictor

Visualization of CpG-age relationships can be viewed using MethylBrowsR: https://shiny.igmm.ed.ac.uk/MethylBrowsR/

## Funding

### Generation Scotland

Generation Scotland received core support from the Chief Scientist Office of the Scottish Government Health Directorates (CZD/16/6) and the Scottish Funding Council (HR03006). Genotyping and DNA methylation profiling of the Generation Scotland samples was carried out by the Genetics Core Laboratory at the Edinburgh Clinical Research Facility, Edinburgh, Scotland and was funded by the Medical Research Council UK and the Wellcome Trust (Wellcome Trust Strategic Award STratifying Resilience and Depression Longitudinally (STRADL; Reference 104036/Z/14/Z). The DNA methylation data assayed for Generation Scotland was partially funded by a 2018 NARSAD Young Investigator Grant from the Brain & Behavior Research Foundation (Ref: 27404; awardee: Dr David M Howard) and by a JMAS SIM fellowship from the Royal College of Physicians of Edinburgh (Awardee: Dr Heather C Whalley).

### Lothian Birth Cohorts

We thank the LBC1921 and LBC1936 participants and team members who contributed to these studies. The LBC1921 was supported by the UK’s Biotechnology and Biological Sciences Research Council (BBSRC), The Royal Society, and The Chief Scientist Office of the Scottish Government. The LBC1936 is supported by the BBSRC, and the Economic and Social Research Council [BB/W008793/1] (which supports S.E.H.), Age UK (Disconnected Mind project), the Medical Research Council (MR/M01311/1), and the University of Edinburgh. Methylation typing of LBC1936 was supported by the Centre for Cognitive Ageing and Cognitive Epidemiology (Pilot Fund award), Age UK, The Wellcome Trust Institutional Strategic Support Fund, The University of Edinburgh, and The University of Queensland. Genotyping was funded by the BBSRC (BB/F019394/1). S.R.C. is supported by a Sir Henry Dale Fellowship jointly funded by the Wellcome Trust and the Royal Society (Grant Number 221890/Z/20/Z).

D.A.G. is supported by funding from the Wellcome Trust 4 year PhD in Translational Neuroscience: training the next generation of basic neuroscientists to embrace clinical research [108890/Z/15/Z]. R.F.H is supported by an MRC IEU Fellowship. M.R.R. was funded by Swiss National Science Foundation Eccellenza Grant PCEGP3-181181 and by core funding from the Institute of Science and Technology Austria. C.H. is supported by an MRC Human Genetics Unit programme grant ‘Quantitative traits in health and disease’ (U. MC_UU_00007/10). E.B. and R.E.M. are supported by Alzheimer’s Society major project grant AS-PG-19b-010.

**This research was funded in whole, or in part, by the Wellcome Trust (104036/Z/14/Z, 108890/Z/15/Z, and 221890/Z/20/Z). For the purpose of open access, the author has applied a CC BY public copyright licence to any Author Accepted Manuscript version arising from this submission**.

