## Supplementary Figures for "Refining epigenetic prediction of chronological and biological age"

**Supplementary Figure 1**. Manhattan plots of (a) linear and (b) quadratic age EWAS in Generation Scotland. Dashed line indicates epigenome-wide significant level of $p = 3.6 \times 10^{-8}$. Genomic position indicated in x-axis, with -log10(*p*-value) of association indicated in y-axis. *p*-values capped at $10^{-320}$.

(a)

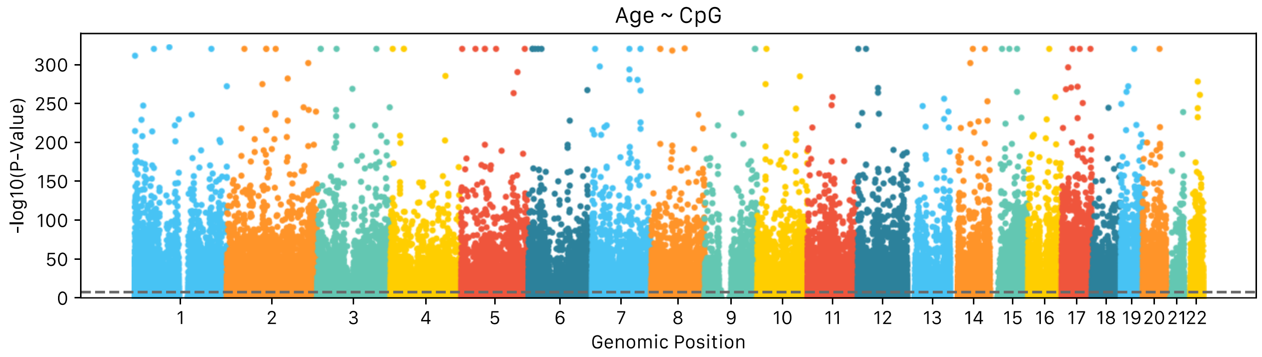


(b)

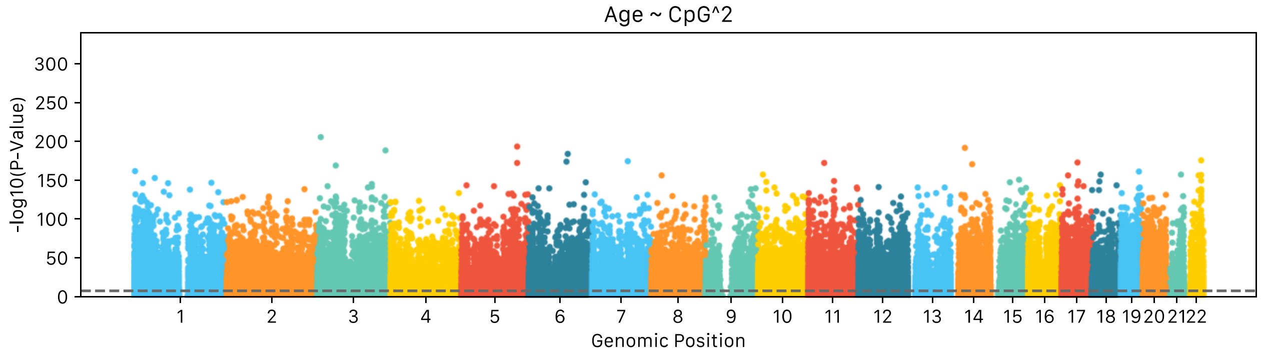

**Supplementary Figure 2**. Scatterplots of top 15 associations from age ~ CpG EWAS. CpG beta values uncorrected for covariates (y-axis) are plotted against age (x-axis) for an unrelated subset of Generation Scotland participants (N = 4,450). LOESS curve with 95% confidence interval and contour plot overlaid. Gene names are taken from the Illumina EPIC array annotation file.

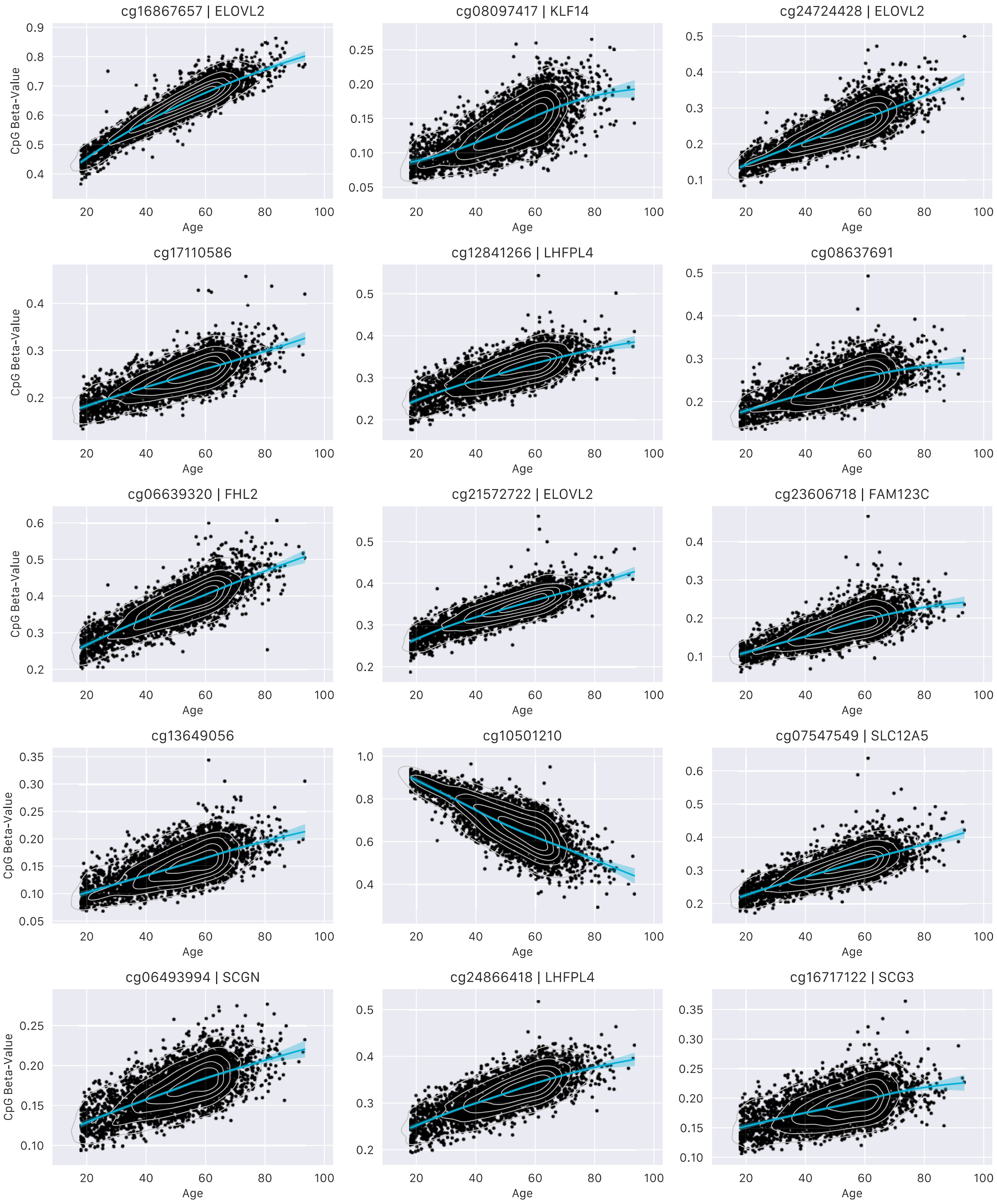

**Supplementary Figure 3**. Scatterplots of top 15 associations from the Age ~ CpG + $CpG^2$ EWAS. Significant associations are based on p-values obtained for the regression coefficient associated to $CpG^2$. CpG beta values uncorrected for covariates (y-axis) are plotted against age (x-axis) for an unrelated subset of Generation Scotland participants (N = 4,450). LOESS curve with 95% confidence interval and contour plot overlaid. Gene names are taken from the Illumina EPIC array annotation file.

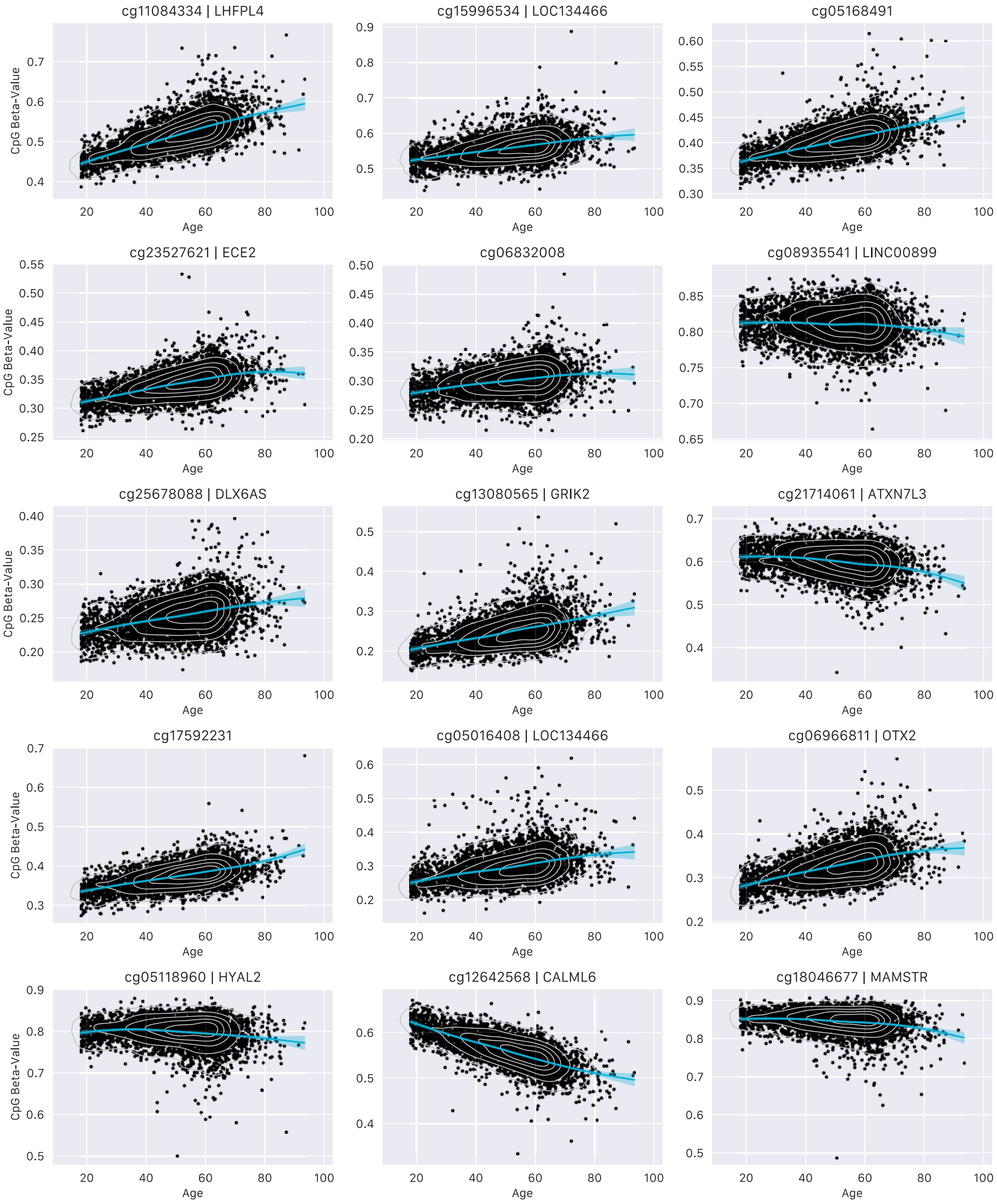

**Supplementary Figure 4**. Prediction metrics (Pearson correlation - r, root mean square error - RMSE, and median absolute error - MAE) for GSE40279, as a function of CpGs included in training of a cAge predictor, trained using Generation Scotland data. CpG subsets obtained from linear age EWAS in Generation Scotland.

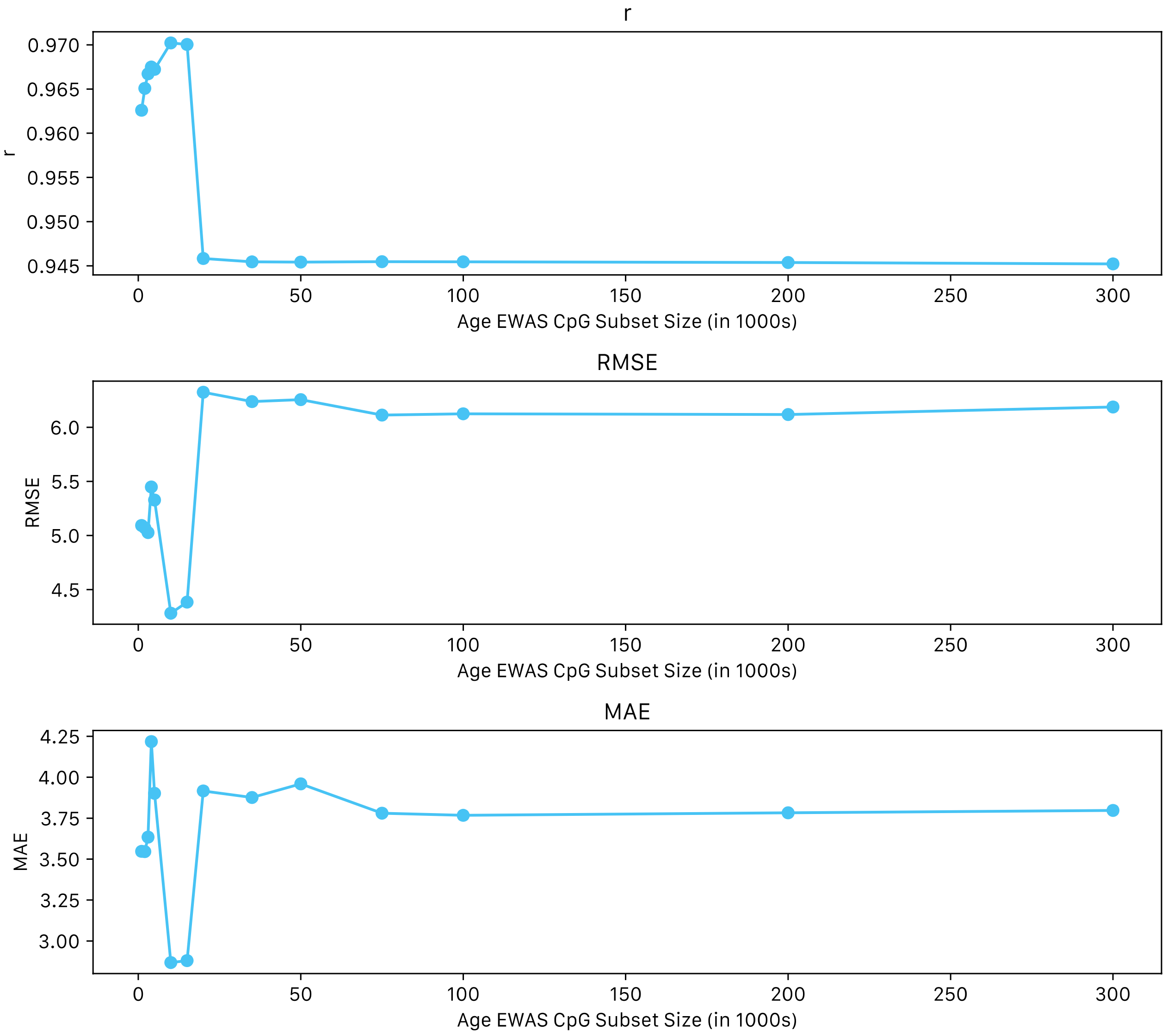

**Supplementary Figure 5**. Prediction metrics (Pearson correlation - r, root mean square error - RMSE, and median absolute error - MAE) for GSE40279, as a function of CpG$^2$s included in training of a cAge predictor trained using Generation Scotland data, in addition to top 10K age-associated CpGs. CpG$^2$ subsets obtained from quadratic age EWAS in Generation Scotland.

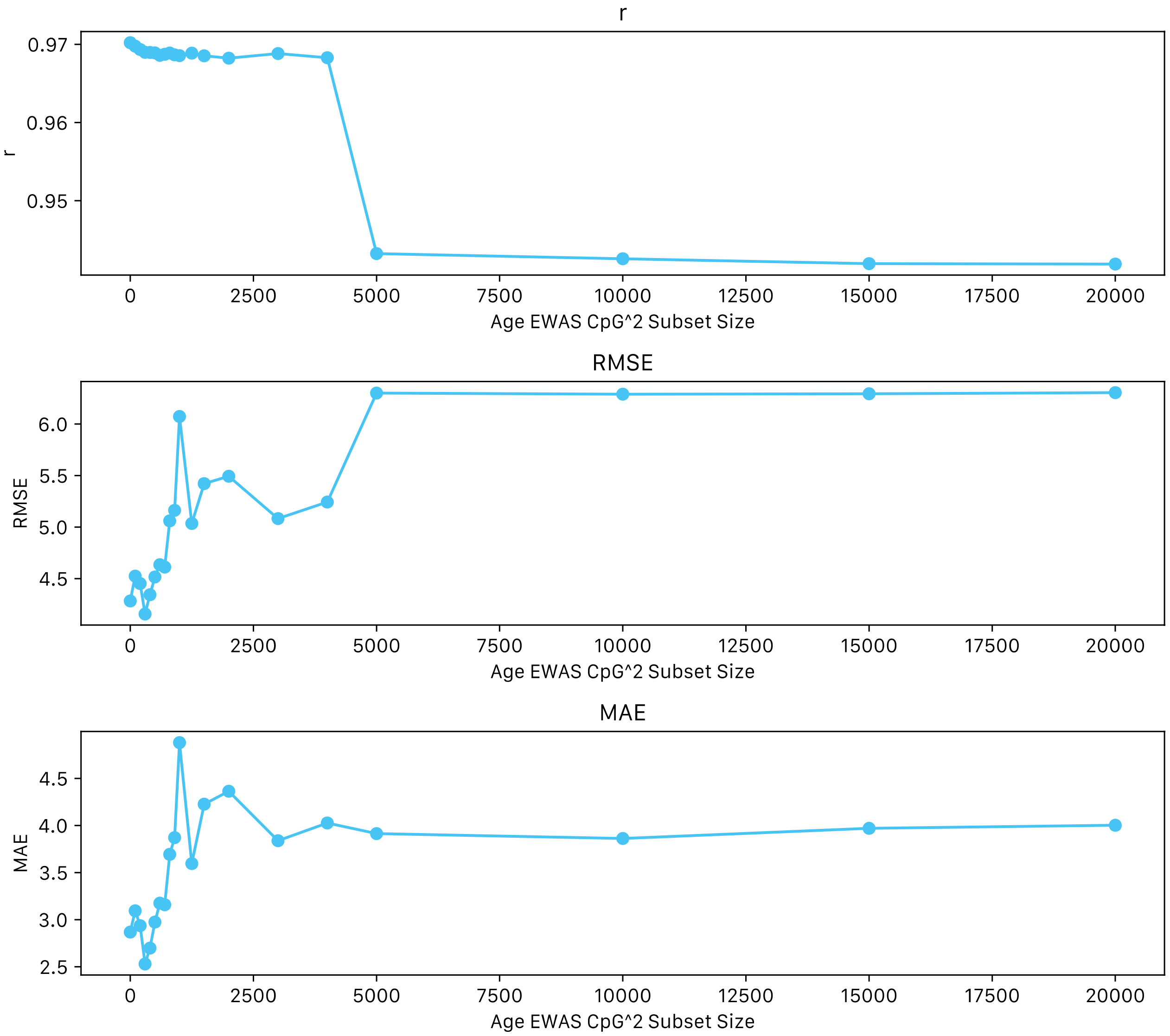

**Supplementary Figure 6**. Manhattan plot of all-cause mortality EWAS in Generation Scotland. Dashed line indicates epigenome-wide significant level of p = 3.6 x $10^{-8}$. Genomic position indicated in x-axis, with -log10(*p*-value) of association indicated in y-axis.

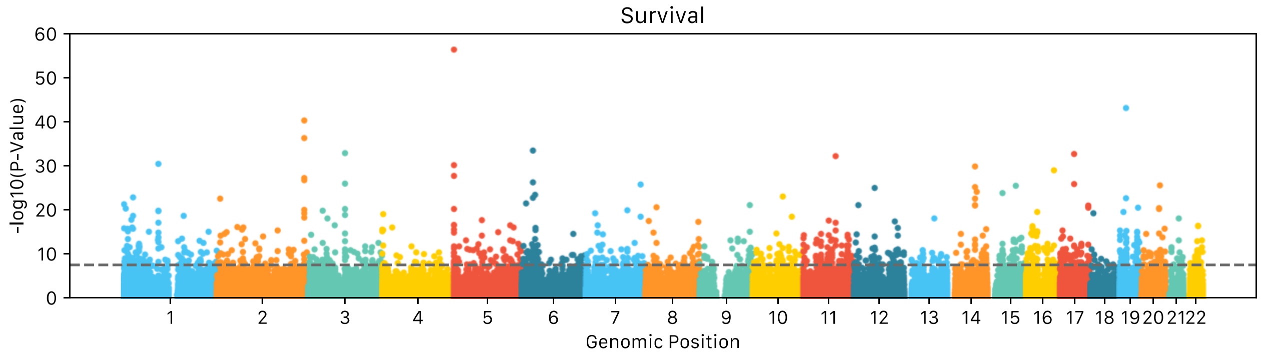

**Supplementary Figure 7**. Comparison of Z-values for 200 overlapping epigenome-wide significant CpG-mortality associations reported by Colicino et al[4], and those considered in the present study (Pearson correlation, r). The grey line shows y = x (perfect correspondence). The correlations for the subset of findings from Colicino et al. with Z>0 and Z<0 are 0.29 and 0.46, respectively.

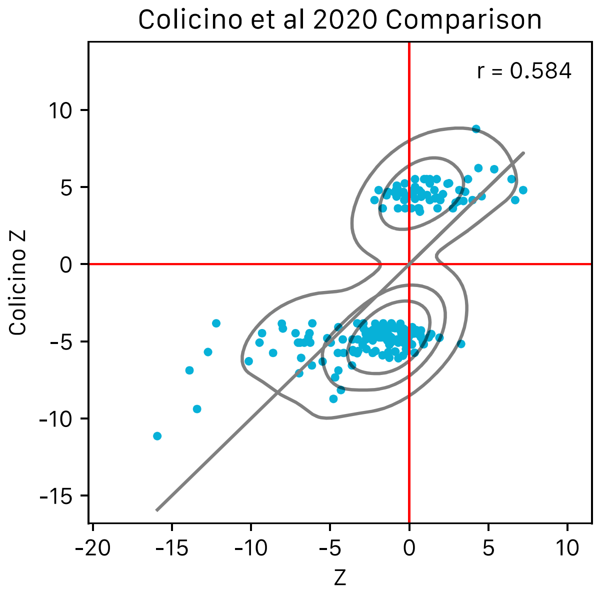
